# Novel small molecule agonists of an *Aedes aegypti* neuropeptide Y receptor block mosquito biting behavior

**DOI:** 10.1101/393793

**Authors:** Laura B. Duvall, Lavoisier Ramos-Espiritu, Kyrollos E. Barsoum, J. Fraser Glickman, Leslie B. Vosshall

## Abstract

Female *Aedes aegypti* mosquitoes bite humans to obtain a blood-meal to develop their eggs. Remarkably, strong attraction to humans is suppressed for several days after the blood-meal by an unknown mechanism. We investigated a role for neuropeptide Y (NPY)-related signaling in this long-term behavioral suppression, and discovered that drugs targeting human NPY receptors modulate mosquito host-seeking behavior. In a screen of all 49 predicted *Ae. aegypti* peptide receptors, we identified NPY-like receptor 7 (NPYLR7) as the sole target of these human drugs. To obtain small molecule agonists selective for NPYLR7, we carried out a high-throughput cell-based assay of 265,211 compounds, and isolated 6 highly selective NPYLR7 agonists that inhibit mosquito attraction to humans. *NPYLR7* CRISPR-Cas9 null mutants are defective in behavioral suppression, and resistant to these drugs. Finally, we show that these drugs are capable of inhibiting biting and blood-feeding on a live host, suggesting a novel approach to control infectious disease transmission by controlling mosquito behavior.

## Introduction

*Ae. aegypti* mosquitoes require blood from a host to complete their full reproductive life cycle. Proteins from blood trigger and sustain egg maturation, and female mosquitoes of this species that fail to obtain a blood-meal do not reproduce. Importantly, a single female will go through multiple blood-feeding and egg-laying cycles in her lifetime. This cycling behavior makes mosquitoes particularly effective disease vectors: when they bite infected humans, disease-causing viral pathogens are passed to the mosquito, and each subsequent bite puts the next human host at risk for infection with dengue, Zika, yellow fever, and chikungunya viruses (Bhatt et al., 2013). Preventing mosquitoes from biting humans is an important point of intervention in global public health strategy.

Although female *Ae. aegypti* mosquitoes are strongly attracted to human hosts when they seek a blood-meal, this attraction is potently inhibited for several days after a complete blood-meal, a period when they digest the blood and mature their eggs (Davis, 1984a, b; Judson, 1967; Klowden, 1990; Klowden and Lea, 1979a, b). Host-seeking suppression consists of at least two phases: a short-term phase likely involving abdominal distension from a blood-meal that doubles the female’s body weight (Klowden and Lea, 1979a), and a long-term sustained phase that lasts until the female lays her eggs (Klowden, 1981; Liesch et al., 2013). The mechanisms that maintain sustained host-seeking suppression following clearance of the blood-meal during egg development are unknown.

Previous studies showed that injection of hemolymph from blood-fed females or high doses of synthetic peptides that activate G protein-coupled neuropeptide Y (NPY)-like receptors were sufficient to suppress host attraction in non-blood-fed females (Brown et al., 1994; Christ et al., 2017; Liesch et al., 2013). These findings suggested that activation of an NPY-related pathway plays a role in sustained host-seeking suppression in *Ae. aegypti* mosquitoes. Early work pointed to Head Peptide-I as the relevant peptide signal mediating suppression (Brown et al., 1994), but more recent studies demonstrated that neither Head Peptide-I nor its receptor NPY-like receptor 1 mediate female host-seeking suppression (Duvall et al., 2017).

Although the exact peptide and receptor combination remains unknown, NPY-related signaling pathways mediated by other peptides and receptors are strong candidates for controlling this behavior because they are key regulators of motivated feeding behavior and satiety across species. NPY-related signaling pathways are deeply evolutionarily conserved and have been implicated in foraging and motivated feeding behavior in organisms including nematodes, mice, rats, and humans (Colmers and Wahlestedt, 1993; de Bono and Bargmann, 1998; Inui, 1999; Maeda et al., 2015; Mekata et al., 2017; Ohno et al., 2017; Wu et al., 2003; Wu et al., 2005). In both vertebrates and invertebrates, there is extensive cross-talk between multiple NPY-like peptides and NPY receptors, with each peptide activating multiple receptors, and a given receptor responding to more than one peptide. For instance, humans have 4 NPY receptors, and their activation has heterogeneous effects on food intake depending upon the specifics of peptide/receptor pairing (Gerald et al., 1996; Larhammar et al., 1992; Lutz et al., 1997; Rose et al., 1995; Tatemoto et al., 1982; Wahlestedt et al., 1986). Invertebrate NPY-like receptors show roughly 60% sequence similarity to vertebrate NPY Y2 receptors (Nässel and Wegener, 2011). Consistent with their evolutionary conservation, small molecule and peptide drugs designed to target human neuropeptide receptors are also capable of acting on insect receptors (de Jong-Brink et al., 2001; Larhammar and Salaneck, 2004).

In this study we show that NPY-like signaling is a central mechanism by which *Ae. aegypti* mosquito attraction to humans is suppressed for several days after a blood-meal. We fed human NPY Y2 receptor drugs to mosquitoes and showed that the drugs interfere with this behavioral suppression. By heterologous expression of all 49 mosquito peptide receptors, we identified a single receptor, NPY-like receptor 7 (NPYLR7), as the sole target of these drugs. We used this cell-based assay to carry out a high-throughput screen of 265,211 compounds to identify 24 new small molecules that activate *Ae. aegypti* NPYLR7 but not human NPY receptors *in vitro*. Of these, 6 have *in vivo* efficacy and suppress mosquito attraction to humans. To demonstrate that NPYLR7 is the sole *in vivo* target of these drugs, we used CRISPR-Cas9 to generate *NPYLR7* mutants. These animals show defects in suppression after a blood-meal, and are resistant to both the human NPY Y2 receptor agonist and our new small molecule NPYLR7 agonists. Finally, we demonstrate that these drugs inhibit host-seeking, biting, and blood-feeding when mosquitoes are offered a live host. These behaviors are the first step in mosquito-borne disease transmission, and exploiting this NPYLR7-dependent endogenous mechanism of host-seeking suppression represents a new strategy for the control of disease-vectoring mosquitoes.

## Results

### Protein-rich blood-meals induce sustained host-seeking suppression

To explore the relationship between dietary composition and host-seeking suppression, we fed female *Ae. aegypti* mosquitoes sheep blood, protein-rich artificial blood (Kogan, 1990), or protein-free saline from a Glytube membrane feeder (Figure 1A) (Costa-da-Silva et al., 2013). Artificial blood is a protein-rich meal containing only purified human albumin, γ-globulins, hemoglobin, and ATP in a sodium bicarbonate solution (Kogan, 1990). The saline meal contains only sodium bicarbonate and ATP, a known feeding stimulant in mosquitoes (Galun et al., 1963). A majority of females fed to repletion on all three meals, as evidenced by an enlarged abdomen and doubling of body weight (Figure 1A-C). Females reliably produced eggs after consuming both protein-rich meals, but never produced eggs after feeding on protein-free saline (Figure 1D). Females produced more eggs with the sheep blood-meal (Figure 1D), likely because of its higher total protein content and complexity of other nutrients, including vitamins, cholesterol, and fatty acids (Kogan, 1990). To ensure that eggs from artificial blood-meals were viable and healthy, we maintained animals exclusively on artificial blood for 4 generations with no loss of viability (data not shown). These experiments confirm that protein is necessary and sufficient for females to develop and lay viable eggs.

**Figure 1.**
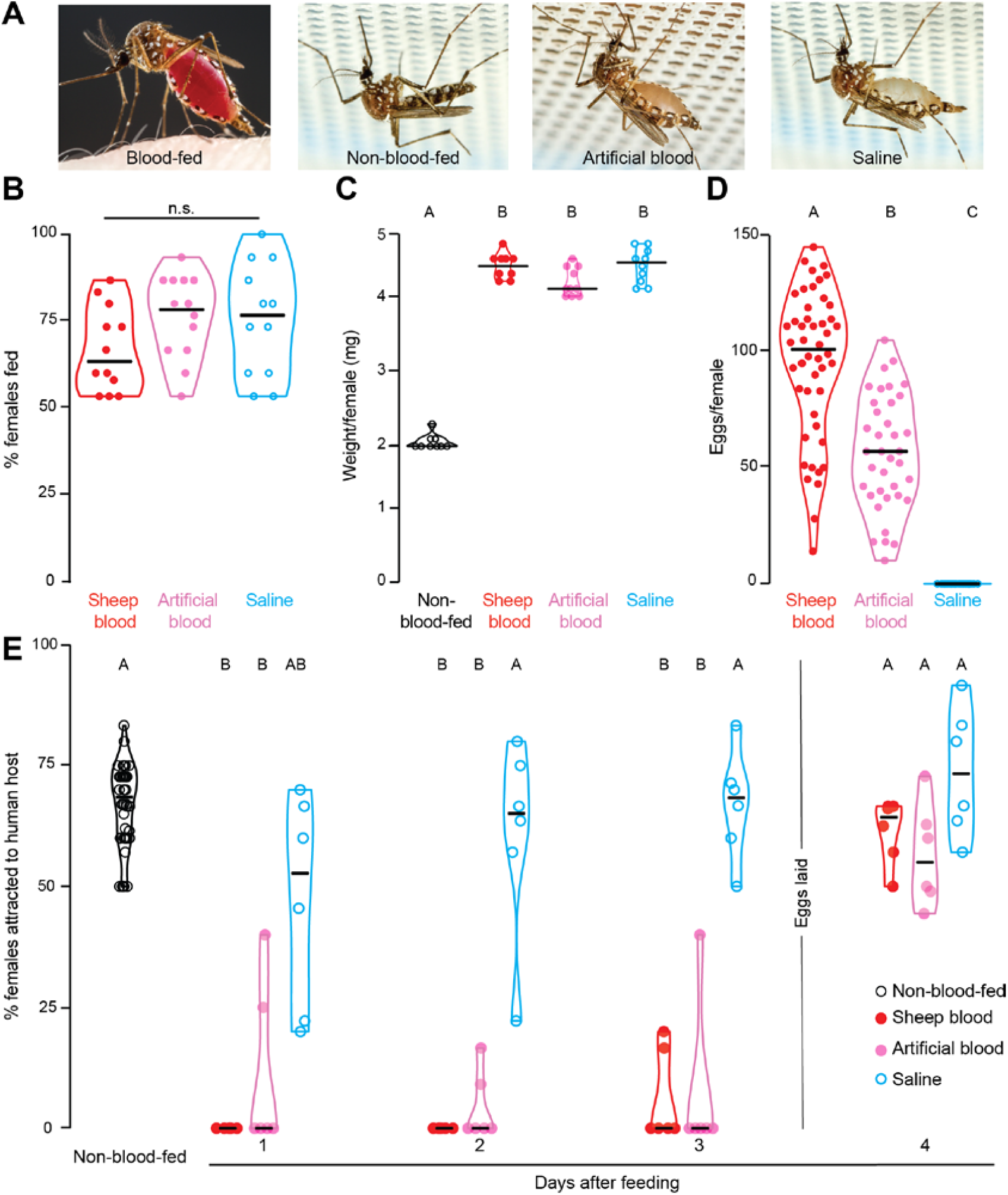
Protein-rich blood-meals induce sustained host-seeking suppression. (A) *Ae. aegypti* females feeding on human skin or Glytube membrane feeders (photos by Alex Wild). (B) Female engorgement on the indicated meal delivered via Glytube (median with range, n=12, 10 - 15 females/trial, Kruskal-Wallis test with Dunn’s multiple comparison, n.s.: not significant p>0.05). (C) Weight per female of the indicated meal (median with range, n=12, 10 - 15 females/trial, one-way ANOVA followed by Tukey’s multiple comparison test). (D) Eggs produced per female after feeding on the indicated meal (median with range, n=37 - 48 females, Kruskal-Wallis test with Dunn’s multiple comparison). (E) Time course of human host-seeking in the uniport olfactometer after feeding with the indicated meal (median with range, n= 6 - 48, 15 - 25 females/trial, Kruskal-Wallis test with Dunn’s multiple comparison). Data labeled with different letters in C-E are significantly different (p<0.05).

To ask how these three different meals affect host-seeking suppression after feeding, we carried out behavioral experiments with a uniport olfactometer (Liesch et al., 2013), which quantifies mosquito attraction to the arm of a live human host and carbon dioxide. Consistent with previous reports, both protein-rich meals induced sustained host-seeking suppression that lasted until eggs were laid, here between days 3 and 4 after feeding (Figure 1E) (Klowden, 1981; Liesch et al., 2013). We note that protein-free saline induced partial short-term host-seeking suppression, likely due to abdominal distension and increased body weight after engorging with this liquid (Klowden and Lea, 1979a). It takes between 1-2 days for females to clear the saline meal, and indeed saline-fed animals returned to normal host-seeking 2 days after feeding (Figure 1E). These findings show that, although female mosquitoes will engorge on both protein-rich and protein-free meals, only protein-rich meals trigger sustained host-seeking suppression and egg production.

### Human NPY receptor drugs modulate mosquito host-seeking behavior

How does blood-feeding inhibit mosquito attraction to humans for multiple days? Prior work in *Ae. aegypti* mosquitoes suggested a role for humoral signals and NPY-like peptides, but the specific molecular mechanism of this inhibition remained unclear. To gain an entry point into the study of peptide signaling in mosquito behavior, we used a pharmacological approach. Because NPY signaling is associated with food intake and obesity in humans, the pharmaceutical industry has developed NPY receptor modulators as therapeutics (Brothers and Wahlestedt, 2010). Inspired by the conserved nature of NPY-like signaling, we reasoned that human NPY receptor drugs might affect cognate receptors in the mosquito and modulate their attraction to humans after a blood-meal. If activation of an NPY pathway mediates host-seeking suppression, human NPY receptor agonists fed in a saline meal should induce suppression, while human NPY receptor antagonists fed in a blood-meal should interfere with suppression.

To test these hypotheses we developed a new, higher-throughput behavioral assay, the miniport olfactometer (Figure 2A), and screened 10 human NPY receptor agonists and antagonists for an effect on behavioral suppression by feeding them to female mosquitoes via a Glytube (Figure 2B). Agonists were fed in non-suppressing saline meals, and antagonists were fed in suppressing artificial blood-meals, and females were assayed for host-seeking in the miniport olfactometer 2 days after drug-feeding. The drugs had no effect on feeding rates or meal size, indicating that they are palatable (Supplemental Data File 1). Of 10 compounds screened, 2 agonists and 1 antagonist affected mosquito host-seeking behavior (Figure 2B and Supplemental Data File 1). TM30335 and TM30338 are structurally similar stabilized peptide mimics of NPY that activate human NPY Y2 and Y4 receptors (Schwartz, 2008). Both drugs induced dose-dependent host-seeking suppression when fed in saline, with TM30335 showing higher potency (Figure 2C and Supplemental Data File 1). In a control experiment, we showed that TM30335 did not affect gross motor behavior (Figure 2D), suggesting a selective effect on suppressing mosquito host-seeking behavior. At the three highest doses, TM30335 was equivalent to a blood-meal in suppressing mosquito attraction to human host cues (Figure 2C). Conversely, BIIE0246, a small molecule antagonist of the human NPY Y2 receptor (Doods et al., 1999), showed dose-dependent release from suppression when fed in a protein-rich artificial blood-meal (Figure 2E). It is notable that all 3 behaviorally active drugs target the NPY Y2 receptor, the human receptor most closely related to insect NPY-like receptors (Brothers and Wahlestedt, 2010; Doods et al., 1999; Kamiji and Inui, 2007; Larhammar and Salaneck, 2004; Nässel and Wegener, 2011). These results suggest that drugs targeting human NPY Y2 receptors are biologically active in mosquitoes, and that endogenous mosquito host-seeking suppression may be mediated in part by NPY-like signaling.

**Figure 2.**
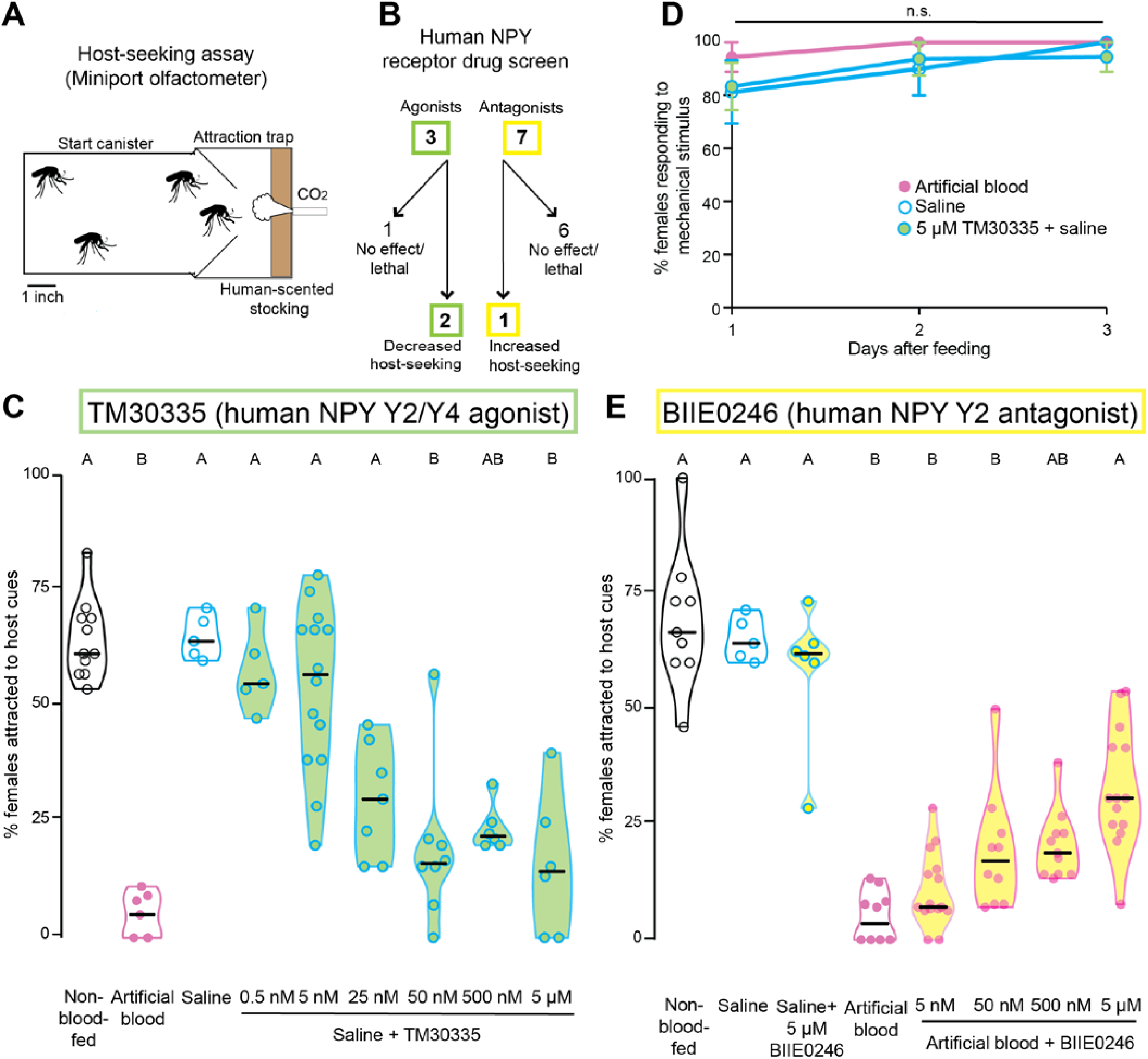
Human NPY receptor drugs modulate *Ae. aegypti* host-seeking behavior. (A) Schematic of miniport olfactometer. Mosquitoes not drawn to scale. (B) Flowchart of human NPY receptor compound behavioral screen. (C) Effect of human NPY Y2/Y4 agonist TM30335 on host-seeking in a miniport olfactometer (n= 5 - 14, 15 - 25 females/trial). (D) Responses to mechanical stimulus in females fed the indicated meals prior to the indicated time (n = 3 - 8, 12 - 16 females/trial, Kruskal-Wallis test with Dunn’s multiple comparison n.s: not significant, p>0.05). (E) Effect of human NPY Y2 antagonist BIIE0246 on host-seeking in a miniport olfactometer (n= 5 – 15, 15 – 25 females/trial). Data in C and E are plotted as median with range and data in D are plotted as median with interquartile range Data labeled with different letters in C and E are significantly different (Kruskal-Wallis test with Dunn’s multiple comparison, p<0.05).

### Among 49 *Ae. aegypti* peptide receptors, NPY-like receptor 7 is the primary target of human NPY receptor drugs

Although the drugs that we tested in mosquitoes are known to target human NPY Y2 receptors, we could not assume *a priori* that they are selectively targeting NPY-like receptors in the mosquito. We therefore cloned all 49 predicted *Ae. aegypti* neuropeptide receptors from both the L3 (Nene et al., 2007) and L5 (Matthews et al., 2017) genome assemblies and geneset annotations, and developed a high-throughput cell-based assay to identify mosquito receptor(s) that are candidate *in vivo* targets of human NPY receptor drugs. Among these are receptors for diverse peptides, and 8 NPY-like receptors (Liesch et al., 2013). We expressed receptor cDNAs in HEK293T cells co-transfected with a promiscuous G protein (Gqα15) (Offermanns and Simon, 1995) and the genetically-encoded calcium sensor GCaMP6s (Chen et al., 2013) in a 384 well plate configuration. Activation of a given receptor is read out by calcium-induced increases in fluorescence of GCaMP6s. Three out of forty-nine receptors tested responded to a 10 μM dose of agonist TM30335 compared to non-transfected controls. Two of these receptors belong to the NPY-like receptor family (NPYLR5 and NPYLR7) (Liesch et al., 2013) (Figure 3A and B).

**Figure 3.**
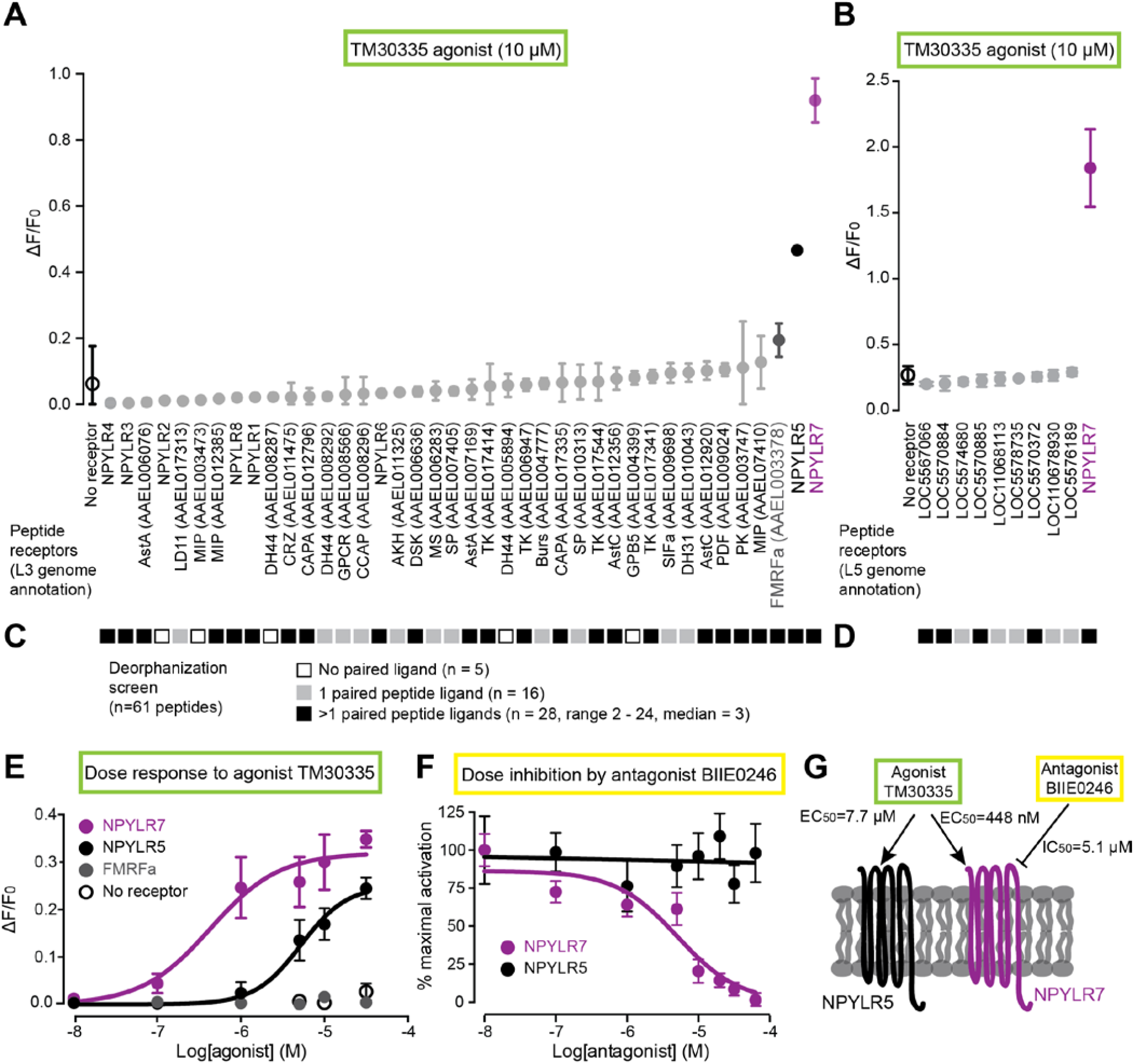
*In vitro* screen identifies NPY-like receptor 7 as the target of behaviorally active compounds. (A-B) Response to TM30335 of all *Ae. aegypti* peptide receptors annotated in the L3 genome assembly and annotation (A) and additional receptors annotated in the L5 genome assembly and annotation (B) (mean ± SD, n=3 trials, 4 replicates/trial). Non-gray data points are statistically different from non-transfected control (black open circle), one-way ANOVA with Bonferroni correction, p<0.001. (C-D) *In vitro* response of 61 predicted *Ae. aegypti* peptides against all predicted peptide receptors in the L3 genome annotation (C) and the L5 genome annotation (D). See Supplemental Data File 1 for raw data. (E) Dose-response curve of TM30335 against the indicated receptors. NPYLR7 EC_50_ = 448 nM, NPYLR5 EC_50_ = 7.7 μM calculated using log(agonist) versus response nonlinear fit (mean ± SD, n=3 trials,4 replicates/trial). (F) Dose-inhibition curve of TM30335 activation NPYLR7 and NPYLR5 responses by BIIE0246. NPYLR7 IC_50_ =5.1 μM calculated using log(inhibitor) versus response nonlinear fit (mean ± SD, n=3 trials, 4 replicates/trial). (G) Summary of *in vitro* findings.

Most of the 49 *Ae. aegypti* peptide receptors have not been matched to cognate peptide ligands. Our cell-based assay offered us the opportunity to attempt a comprehensive deorphanization of all *Ae. aegypti* peptide receptors. This experiment would reveal the mosquito ligands for NPYLR7, and would also allow us to rule out the possibility that failure of TM30335 to activate some or all of the remaining 46 receptors was due to issues with heterologous expression. To carry out this deorphanization screen, we synthesized peptides corresponding to all 61 detected peptides in *Ae. aegypti* (Predel et al., 2010) (Supplemental Data File 1) and screened them against all 49 predicted peptide receptors. All but 5 receptors showed responses to at least one peptide at a 10 μM dose (Figure 3C and D and Supplemental Data File 1). Because most of the peptide receptors not activated by TM30335 were activated by one or more mosquito peptides, we can exclude the possibility that these receptors did not functionally express in heterologous cells. Further, this confirms that TM30335 has narrow selectivity for a small number of mosquito peptide receptors. Deorphanization of the remaining 44 receptors revealed a range of specificities for peptide ligands. 16 receptors were activated by only a single peptide ligand, while the other 28 receptors were activated by 2 or more peptides. The FMRFa receptor showed very broad ligand tuning, with 24 of 61 peptides activating this receptor. NPYLR7 was more selective, with 9 peptide ligands (FMRFa1, FMRFa 3, FMRFa10, MIP-1, sNPF-1, sNPF2 +4, sNPF3, leucokinin, and AAEL011702A) (Supplemental Data File 1).

Dose-response analysis showed that while both NPYLR5 and NPYLR7 are activated by TM30335, NPYLR7 is more sensitive to the agonist with an EC_50_ of 448 nM compared to the NPYLR5 EC_50_ of 7.7 μM (Figure 3E). The third receptor from the initial screen, FMRFa receptor, did not respond to TM30335 upon rescreening, suggesting that it was a false positive (Figure 3E). We next asked if the behaviorally active antagonist BIIE0246 acts on the same receptors activated by TM30335. Dose-inhibition analysis showed that BIIE0246 inhibited TM30335 activation of NPYLR7 with an IC50 of 5.1 μM, but had no effect on NPYLR5 responses (Figure 3F). Since NPYLR7 is the primary peptide receptor targeted by the human drugs that modulate mosquito behavior, it is a strong candidate for mediating host-seeking suppression *in vivo* (Figure 3G).

### *NPYLR7* mutants fail to sustain host-seeking suppression

To test the role of NPYLR7 in host-seeking regulation, we used CRISPR-Cas9 gene editing (Kistler et al., 2015) to generate a null mutation in this gene. We obtained a 4 basepair deletion in the second exon of *NPYLR7*, which is predicted to truncate the receptor near the first transmembrane domain (Figure 4A). These mutant animals showed no gross developmental or anatomical abnormalities and blood-fed normally (Figure 4B and C). *NPYLR7* mutant females showed robust host-seeking that was indistinguishable from that of wild-type and heterozygous controls (Figure 4D). However, while the mutants showed normal short-term suppression seen 1 day after the blood-meal, they failed to maintain long-term host-seeking suppression (Figure 4D). These results provide genetic evidence that *NPYLR7* regulates endogenous host-seeking suppression following a blood-meal.

**Figure 4.**
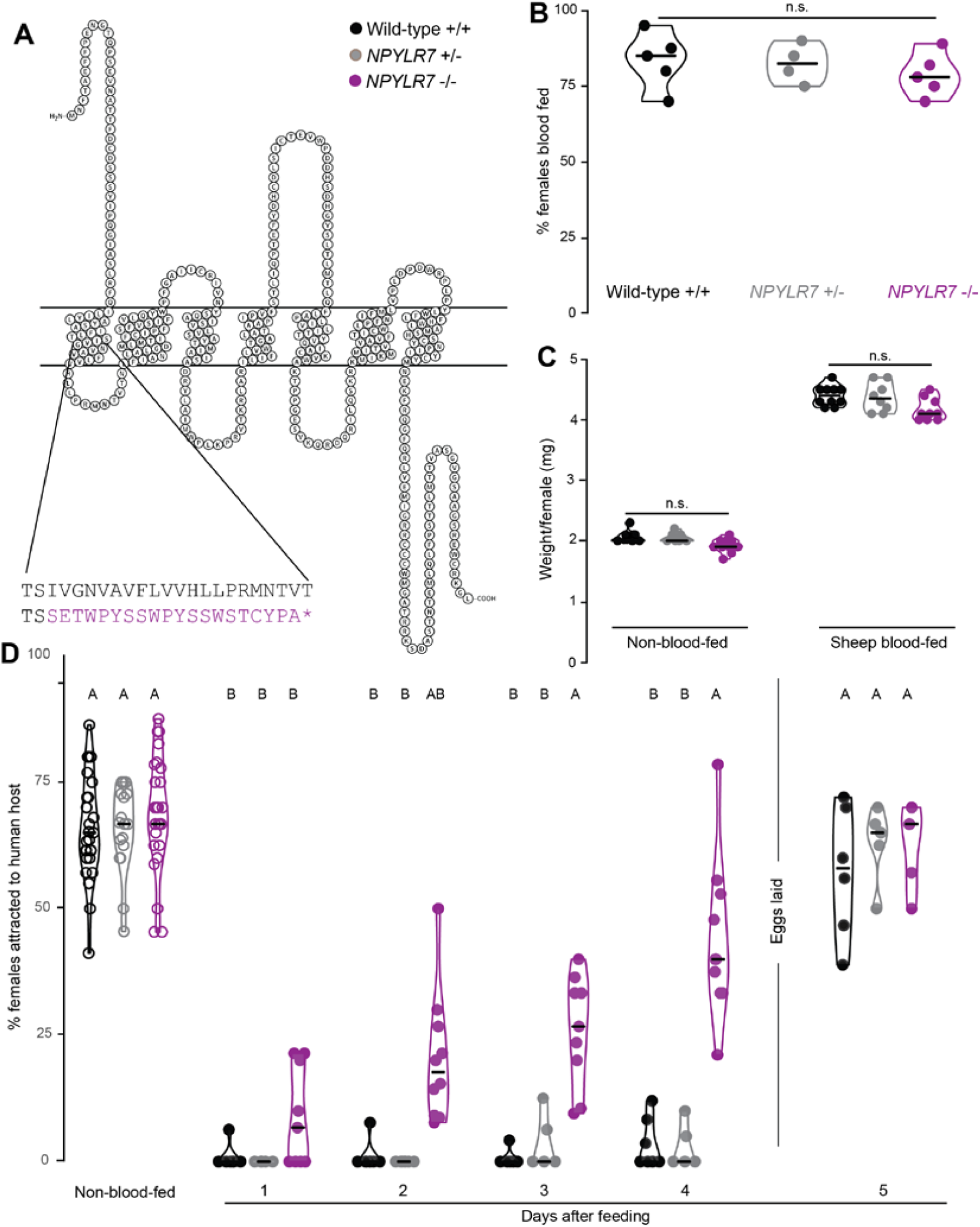
*NPYLR7* mutants blood-feed normally but do not maintain sustained host-seeking suppression. (A) Snake plot of wild-type NPYLR7 and predicted truncated amino acid sequence of *NPYLR7Δ4* mutant. Stop codon is indicated by the purple asterisk. (B) Engorgement of females of the indicated genotype on sheep blood delivered via Glytube (median with range, n=4 – 5, 35 - 50 females/trial, Kruskal-Wallis test with Dunn’s multiple comparison test, n.s., not significant, p>0.05). (C) Weight per female of non-blood-fed and sheep-blood-fed females (median with range, n= 8-10, 10 females/trial, one-way ANOVA with Sidak’s multiple comparisons test, n.s., not significant, p>0.05). (D) Percentage of females attracted to human host in the uniport olfactometer. Females were allowed to lay eggs between days 4 and 5. (median with range, n=5-27, 15-25 females/trial). Data labeled with different letters are significantly different (Kruskal-Wallis test with Dunn’s multiple comparison, p<0.05).

### Small molecule screen identifies NPYLR7 agonists that suppress host-seeking *in vivo*

Triggering endogenous host-seeking suppression via NPYLR7 could be a route to prevent mosquito biting behavior. Although TM30335 is a potent agonist of mosquito NPYLR7, it also activates human NPY receptors. These off-target effects could limit any practical applications of this drug as a mosquito behavioral control agent. With this in mind, we sought to identify more selective small molecule agonists of NPYLR7 that do not activate human NPY receptors. We carried out a high-throughput screen of small molecules to find selective *Ae. aegypti* NPYLR7 agonists (Figure 5A). We screened 265,211 unique small molecules at a concentration of 10 μM using GCaMP6s activation as a readout of NPYLR7 activation, and identified 376 compounds that activated cells expressing NPYLR7. These were then counter-screened against non-receptor transfected controls, in an independent cell-based Fura2 assay, and validated by mass spectrometry (Supplemental Data File 1). This secondary screen identified 24 compounds that specifically and robustly activate NPYLR7 *in vitro* (Figure 5A).

**Figure 5.**
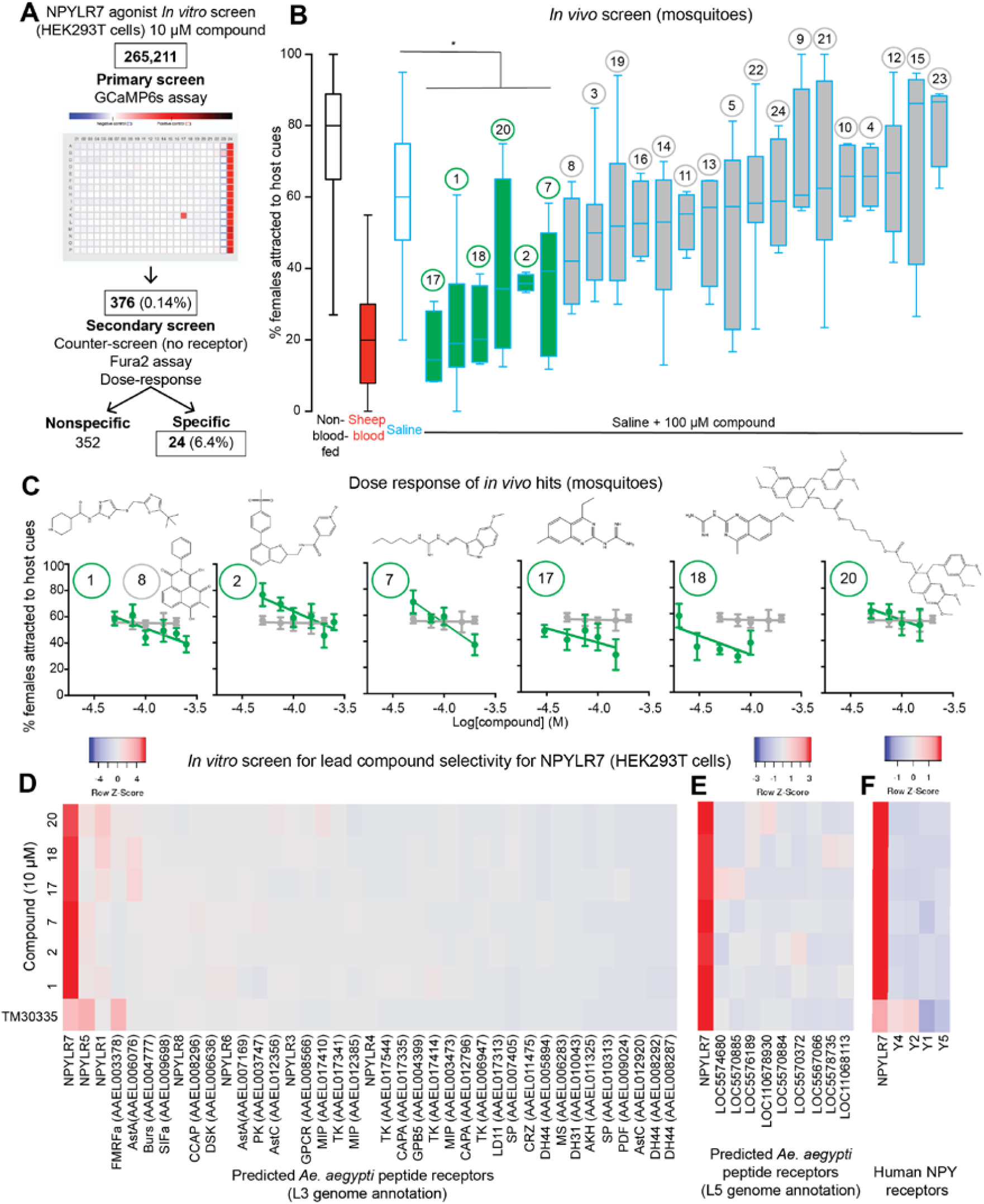
Small molecule screen identifies NPYLR7 agonists with *in vivo* activity. (A) Schematic of high-throughput small molecule screen for NPYLR7 agonists. (B) 24 confirmed *in vitro* hits tested for *in vivo* activity using the miniport olfactometer (median with interquartile range, n = 4 – 116, 15 - 25 females/trial). Compounds are indicated at the top of the figure with identifier number in a circle. Groups in green are significantly different compared to saline meal control (Kruskal-Wallis test with Dunn’s multiple comparison p<0.05). (C) *In vivo* dose-response tests of 6 primary hits in the miniport olfactometer (median with interquartile range, n=4 – 6, 15 - 25 females/trial). Data from inactive compound 8 are replotted in all 6 panels. (D-E) *In vitro* response profile of NPYLR7-activating compounds against all predicted peptide receptors in the L3 (D) and L5 (E) annotation of the *Ae. aegypti* genome. (F) *In vitro* response profile of NPYLR7-activating compounds against human NPY receptors.

To ask if these 24 small molecules could suppress host-seeking behavior *in vivo*, we fed mosquitoes individual compounds in protein-free saline and assayed host-seeking in the miniport olfactometer 2 days after feeding (Figure 5B). Addition of the compounds to saline had no effect on feeding rates or meal size, indicating that all compounds were palatable (Supplemental Data File 1). Compound 6 resulted in >50% lethality and was therefore excluded from further analysis. This behavioral screen identified six small molecule NPYLR7 agonists that suppressed host-seeking in the miniport olfactometer assay (Figure 5B). We then tested lower doses of these hits and determined that compound 18 had the highest potency *in vivo* (Figure 5C). These data show that the compounds identified in our cell-based screen for NPYLR7 agonists are active *in vivo* and are capable of suppressing host-seeking behavior.

To determine the specificity of these six compounds for NPYLR7, we screened them *in vitro* against the panel of all 49 known *Ae. aegypti* peptide receptors from both L3 (Figure 5D) and L5 genome annotations (Figure 5E), as well as human NPY receptors (Figure 5F). All 6 compounds showed high specificity for NPYLR7 *in vitro* with minimal off-target effects on other mosquito peptide receptors. As expected, TM30335 activated human NPY Y2 and Y4 receptors, but the small molecule agonists of NPYLR7 identified in our screen did not activate any human NPY receptors (Figure 5F). We conclude that the hits from our small molecule screen are highly selective NPYLR7 agonists that are capable of inhibiting mosquito attraction to human host cues.

We noted that the two most potent compounds, 17 and 18, were variations of guanidinium-substituted quinazolines, and were the most structurally similar among all of the small molecules in our primary screen. To ask if structurally related compounds were also biologically active, we identified commercially available compounds with quinazoline scaffolds similar to compound 18 (Figure 6A), and tested these analogues in our *in vitro* assay to explore the relationship between structure and activity compared to the initial hit. Compound 18F, which lacks the guanidinium group, showed similar *in vitro* (Figure 6B) and *in vivo* (Figure 6C) potency to the original hit, compound 18. In contrast, compound 18C, with an extended guanidinium group, showed limited *in vitro* efficacy despite structural similarity to active compounds 18 and 18F (Figure 6B) and showed similarly limited *in vivo* efficacy (Figure 6C). Thus, the guanidinium group was not essential for the activity. These results indicate that our *in vitro* assay accurately identifies compounds that activate NPYLR7 *in vitro* and are capable of suppressing host-seeking *in vivo*. The assaying of structurally related compounds demonstrates an initial structure-activity relationship and provides the preliminary characterization of a minimal functional scaffold required for biological activity.

**Figure 6.**
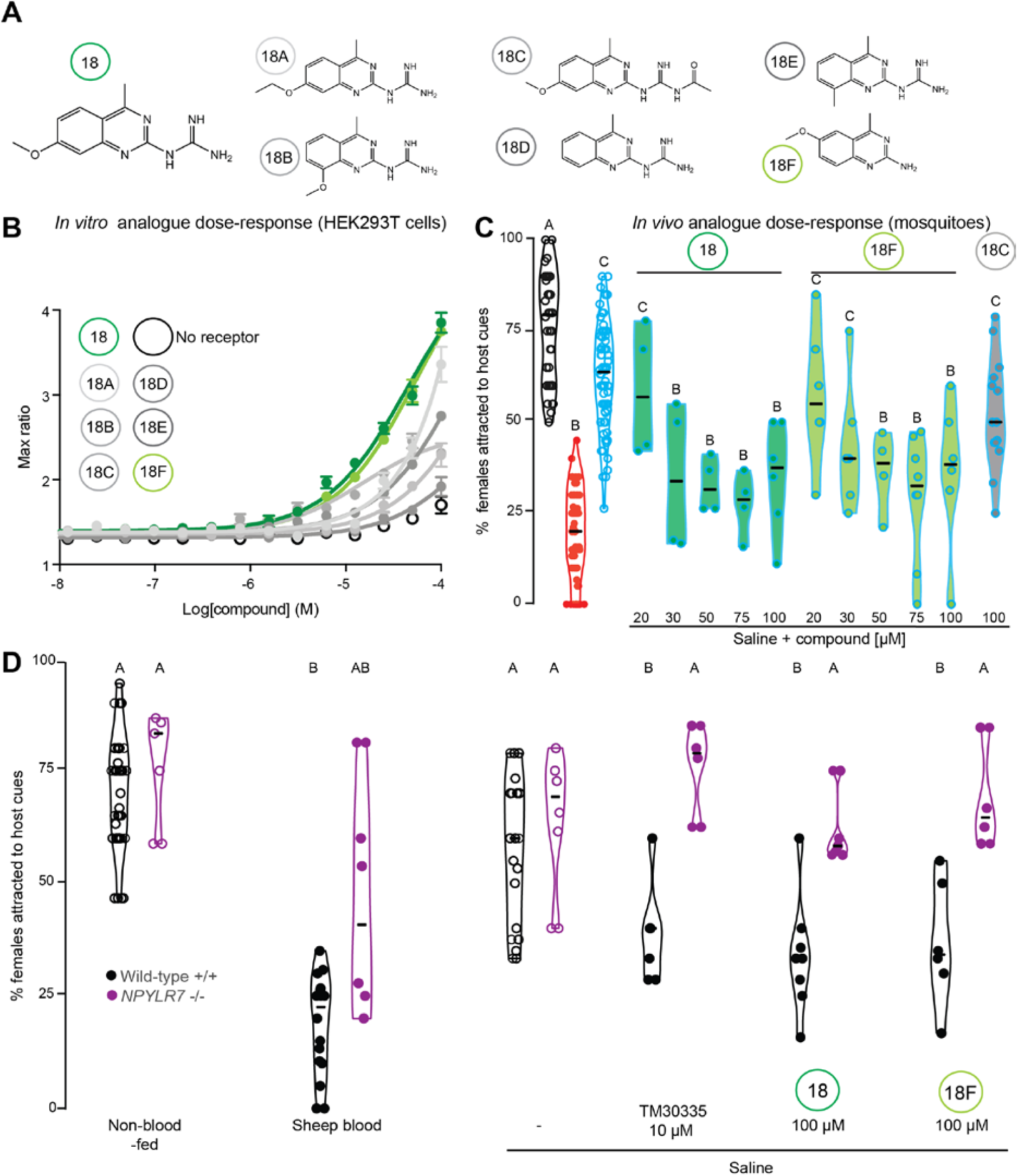
Small molecule screen compounds target *Ae. aegypti* NPYLR7 with high specificity *in vitro* and *in vivo*. (A) Chemical structures of compound 18 and six structural analogues. (B) *In vitro* dose-response curve of compounds in (A) against NPYLR7 (mean ± SEM, n=1, 3 replicates/trial) (C) Host-seeking in a miniport olfactometer 2 days after feeding the indicated meals (n = 4 – 82, 15 - 25 females/trial) (D) Host-seeking in a miniport olfactometer 2 days after feeding the indicated genotypes with the indicated meals (n = 4-26, 15 - 25 females/trial). Data in C-D are plotted as median with range. Data labeled with different letters are significantly different (Kruskal-Wallis test with Dunn’s multiple comparison, p<0.05).

If the behavioral effects of these compounds are mediated exclusively by NPYLR7, we would expect mosquitoes mutant for this receptor to be resistant to drug-induced host-seeking suppression. Consistent with our previous results in the uniport olfactometer, *NPYLR7* mutant females were capable of host-seeking in the miniport olfactometer when non-blood-fed and when fed saline, but showed deficits in sustained host-seeking suppression after a sheep blood-meal (Figure 6D). Consistent with the hypothesis that NPYLR7 is the sole *in vivo* target of the active compounds, *NPYLR7* mutant females were completely resistant to TM30335 as well as compounds 18 and 18F (Figure 6D). This experiment also demonstrates that the drugs do not have side effects that affect mosquito host-seeking behavior beyond their action on NPYLR7.

### NPYLR7 agonists prevent blood-feeding on live hosts in mark-release-recapture experiments

In our previous experiments, we used a miniport olfactometer to demonstrate *in vivo* efficacy of the human NPY receptor drug TM30335 and our new small molecules hits. These assays measure mosquito attraction to human host cues such as odor and carbon dioxide, but do not give mosquitoes access to bite and blood-feed from a live host. Since biting and blood-feeding are the key behaviors that lead to disease transmission, we asked if NPYLR7 agonist compound 18 is capable of preventing mosquitoes from approaching, biting, and blood-feeding on a live host. Female mosquitoes were fed either saline alone, or saline with 100 μM of active compound (18) or inactive compound (18C) and allowed to recover for 2 days. Females from each feeding group were separately marked with one of three different paint powders using the “shake in a bag” technique (Verhulst et al., 2013), and all three of these groups were pooled and allowed to recover together in a cage. Powder marking was randomized between trials to ensure that each drug treatment/powder marked pairing was used and that powder marking did not directly interfere with the ability host-seek, bite, or blood-feed from a live host. After recovery, an anesthetized mouse was placed in the cage and female mosquitoes were given 15 minutes of free access to the mouse before being removed from the cage and scored for blood-feeding (Figure 7A). Females were scored for powder color to indicate treatment condition and for the presence of fresh blood in the midgut (Figure 7A). Similar to our results in host-seeking assays, non-blood-fed and saline-fed females consumed fresh blood at high levels. In contrast, females that had fed on sheep blood 2 days prior, rarely consumed fresh blood. Females fed active compound 18 also showed very low levels of blood-feeding on a live host, comparable to that seen in females suppressed by a prior blood-meal (Figure 7B). Finally, mosquitoes fed inactive compound 18C consumed fresh blood at levels comparable to non-fed females or females fed saline. These experiments demonstrate that this selective NPYLR7 agonist is capable of suppressing host-seeking, biting, and blood-feeding on a live host.

**Figure 7.**
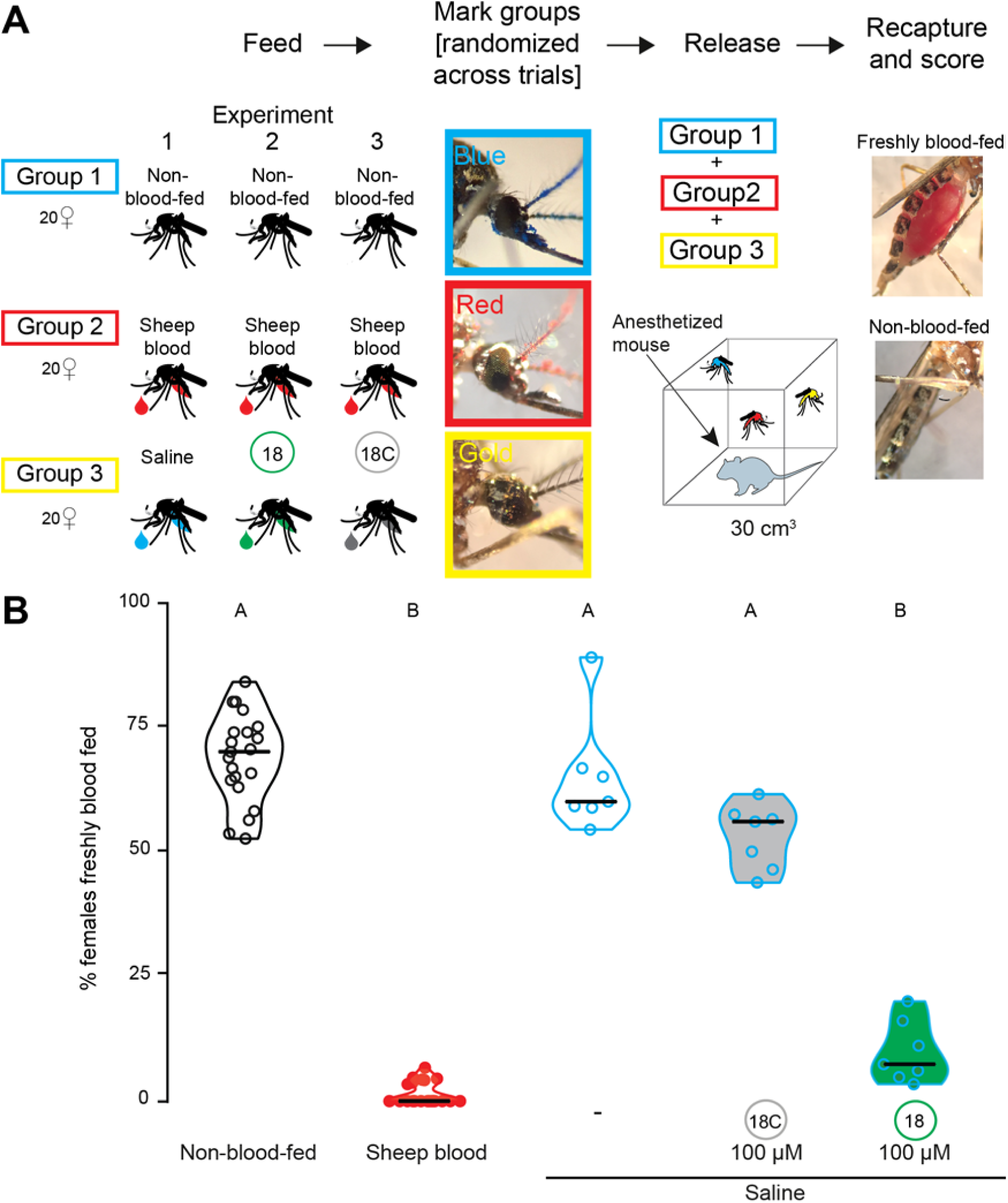
NPYLR7 agonist compound 18 inhibits mosquito blood-feeding on a live host. (A) Schematic of live host assay experiment. (B) Percentage of females, fed the indicated meal 2 days prior to the experiment, which freshly blood-fed on an anesthetized mouse after a 15 min exposure (median with range, n = 7 – 21, 58 - 62 females/trial. Data labeled with different letters are significantly different (Kruskal-Wallis test with Dunn’s multiple comparison, p<0.05).

## Discussion

In this study we identify NPYLR7 as a key member of a peptide signaling pathway that regulates endogenous host-seeking suppression in *Ae. aegypti* mosquitoes. This receptor belongs to the NPY receptor family, which plays a role in feeding behavior across many organisms. We show that both genetic and pharmacological disruption of NPYLR7 signaling leads to abnormal host-seeking regulation, and identify highly specific small molecule agonists of NPYLR7 that can prevent mosquitoes from host-seeking, biting, and blood-feeding.

NPY signaling is deeply conserved and mediates hunger/satiety in organisms ranging from nematodes to humans. Interestingly, NPY signaling can either induce or suppress feeding depending on the ligand/receptor combination. In humans and rodents, activation of NPY Y2 receptor, the most similar to invertebrate NPY-like receptors, reduces food intake (Batterham et al., 2002; Degen et al., 2005). In *Drosophila*, NPY-like signaling pathways, including those mediated by NPF, sNPF, and RYamides peptides, have both positive and negative effects on feeding, metabolism, and growth although their roles are likely pleiotropic, and non-redundant (Nässel and Wegener, 2011; Wu et al., 2003; Wu et al., 2005). Our findings reveal NPY-related signaling in a unique form of motivated feeding behavior in the context of an important disease vector. We propose that after blood-feeding, NPY-like receptor activation leads to sustained host-seeking suppression, which is analogous to a form of long-term satiety. Although our data support a key role for NPYLR7 in mediating host-seeking suppression following a blood-meal, there are clearly also NPYLR7-independent components. Early host-seeking suppression is unrelated to the protein content of the meal, and previous work suggests that abdominal distension regulates this early phase (Klowden and Lea, 1979a). Indeed, the residual host-seeking suppression phenotype of *NPYLR7* mutant females at 2 and 3 days after the blood-meal may be a consequence of slower blood-meal digestion (Supplemental Data File 1). We speculate that mutant females experience a longer period of abdominal distension that may contribute to short-term host-seeking suppression. Perhaps related to delayed digestion, female mutants laid fewer eggs compared to wild-type and heterozygous females (Supplemental Data File 1). Signals from developing eggs, ovary, and fat body have been shown suppress host-seeking, although their molecular identity is unknown (Klowden, 1981; Klowden et al., 1987). Understanding the interactions between these processes and how they combine to produce the complete expression of endogenous host-seeking suppression following a blood-meal remains an important area for future work.

Our work has identified a receptor necessary for host-seeking suppression and exogenous drugs that can activate it, but the endogenous ligand(s) remain unknown. Previous work showed that direct injection of high doses of peptides including sNPF2, sNPF3, and HP-I are capable of suppressing host-seeking behavior (Brown et al., 1994; Christ et al., 2017; Liesch et al., 2013). These peptides all activate NPYLR7 *in vitro* and are likely to exert their effects, at least in part, through NPYLR7 activation (Liesch et al., 2013). In our comprehensive deorphanization of all *Ae. aegypti* peptide receptors, we found additional peptide ligands for NPYLR7 including FMRFamides and leucokinin. Identifying the specific endogenous ligand(s) that mediate host-seeking suppression after a blood-meal remains an important and ongoing area of research, which we expect to be challenging because of extensive redundancy in peptide/receptor interactions.

The identity of the anatomical circuits involved and the mechanisms by which NPY-like signaling interacts with them remain unknown and is currently limited by a lack of genetic reagents in *Ae. aegypti*. Identifying the tissues and cells expressing NPYLR7 is an important goal. Previous work suggested that peptides may be released from midgut endocrine cells that monitor the nutritional content of the midgut, and that their activation leads to peptide release into the hemolymph where these peptides may circulate to act on distant receptor-expressing tissues (Huang et al., 2011). It is possible that blood-meal digestion triggers local release of peptides that activate NPYLR7 directly in the midgut. Alternately, endogenous ligand could be released from neurosecretory cells in the brain in response to blood digestion or egg development signals to activate NPYLR7 either locally in the brain or in visceral organs. RNA-seq data suggest that NPYLR7-expressing cells are found in the female abdominal tip and antennae (Matthews et al., 2016), although it is also possible that this receptor is expressed by a small number of cells in the brain or in the midgut that may not have been detectable. The recent development of genetic tools in the mosquito (Riabinina et al., 2016) will enable future studies of the anatomical and neural circuit basis of host-seeking regulation.

Previous studies suggest that peripheral sensitivity to host-associated cues is down-regulated during host-seeking suppression in *Ae. aegypti* (Siju et al., 2010). It is possible that NPYLR7 regulation acts as a gain control mechanism to alter the peripheral sensitivity to host-associated cues. Work in the *Drosophila* olfactory system has shown that hunger upregulates NPY receptors to increase sensitivity to odors (Root et al., 2011). However, this is likely to be distinct from the pathway described here because NPYLR7 activation suppresses responsiveness to host-associated cues, effectively suppressing hunger and acting as a long-term satiety signal. Our finding that *NPYLR7* mutant females show delayed blood-meal processing also suggests that this pathway may play a role in other aspects of appropriate blood-meal processing including late-stage blood-meal digestion (Gulia-Nuss et al., 2011). Blood-meal digestion, nutrient utilization, and egg development are all key components of host-seeking suppression and the relationships between these processes remain to be completely understood.

We have identified small molecules that suppress mosquito biting when fed at micromolar doses. The compounds identified in our screen are very selective agonists of NPYLR7 with minimal activation of other *Ae. aegypti* receptors or human NPY receptors. Characterization of structure-activity relationships of these compounds will refine these chemical structures to optimize their potency *in vivo*. Our compounds are structurally related to other compounds that act as ligands to human NPFF receptors (Kawakami et al., 2003; Mankus and McCurdy, 2012), yet are notable for their selective activation of insect and not human NPY receptors. Although we have not profiled the *in vivo* half-life of the compounds identified in our small molecule screen, animals maintained suppression for at least 2 days after drug-feeding. This suggests that either the compounds remain active *in vivo* for at least 2 days, are slowly released from the midgut, or that acute NPYLR7 activation leads to downstream signaling pathways that “lock” females into a state of host-seeking suppression for days after drug dosing.

Our results show that feeding small molecule NPYLR7 agonists to female mosquitoes is sufficient to prevent host-seeking, biting, and blood-feeding behavior, raising the possibility that field application of these compounds could reduce disease transmission by inhibiting the drive of mosquitoes to seek humans. These compounds could be delivered in baited attraction traps using stimuli that mimic host-associated cues including human body odor and carbon dioxide (Mathew et al., 2013). Delivering these compounds in traps baited with human odor will also ensure that beneficial insects are not targeted by these methods. Non-lethal methods of pest control have succeeded with other species. Development of a non-toxic compound that destroys immature ovarian follicles in female rats, rendering them sterile has recently shown success and been developed for commercial use (Witmer et al., 2017). Field-based pheromone dispensers have been effective in luring male European grapevine moths away from females to decrease the number of successful matings and thus pest population size (Lucchi et al., 2018).

Although baited traps are often used to deliver toxic compounds, the use of insecticides results in a strong selection pressure on populations by selecting for rare individuals carrying resistance alleles that can escape lethality (Liu, 2015). The widespread deployment of toxic glucose-baited Combat cockroach traps resulted in the rapid selection for an aversion to glucose in populations of German cockroaches (Silverman and Ross, 1994). Unlike cockroaches, which are omnivorous scavengers and can tolerate a loss of glucose preference, mosquitoes are specialized to blood-feed on mammalian hosts and are unlikely become resistant to traps that exploit their attraction to host-associated cues. Additionally, because our compounds target NPYLR7, which is required for appropriate host-seeking suppression, we speculate that females who “escape” the effects of these compounds through natural mutations in *NPYLR7* will have reduced fitness due to the reduced fecundity associated with *NPYLR7* loss of function.

The pattern of active host-seeking followed by host-seeking suppression after a blood meal is observed widely in blood-feeding arthropods including other mosquito species and ticks (Anderson and Magnarelli, 2008; Klowden and Briegel, 1994; Takken et al., 2001), and members of the NPY-like receptor family are found in the genomes of these organisms (Garczynski et al., 2005, 2007; Gulia-Nuss et al., 2016). It is likely that the compounds identified in our screen will show efficacy in additional disease vectors. Exploiting the female mosquito’s endogenous regulation of her own host-seeking behavior represents a novel strategy to prevent the spread of vector-borne disease and may have broad applications across blood-feeding arthropods that spread disease to hundreds of millions of people each year.

## SUPPLEMENTAL INFORMATION

All raw data in the paper are provided in Supplemental Data File 1.

## ACKNOWLEDGMENTS

We thank Emily Dennis, Kevin Lee, Nilay Yapici, and members of the Vosshall Lab for comments on the manuscripts; Alison Ehrlich for technical assistance in receptor cloning, assay development, and strain maintenance; Grace Matthews for assay development while a high school student in the Rockefeller University Summer Science Research Program; Gloria Gordon and Libby Mejia for expert mosquito rearing; Henrik Molina of the Rockefeller University Proteomics Resource Center for proteomic data on endogenous peptides in mosquitoes; and Jim Petrillo of the Rockefeller University Precision Fabrication Facility for assistance with the design and construction of the miniport olfactometer. We thank Vanessa Ruta and members of the Ruta lab for providing guidance and technical assistance in the development of our GCaMP cell-based assays. We thank Thue W. Schwartz (formerly of 7TM Pharma A/S) and Nancy Thornberry and Doug MacNeil (formerly of Merck Research Laboratories) for providing drugs used in this paper. Support for this project was provided by an Advanced Grant from the Robertson Therapeutic Development Fund, generously provided by the Robertson Foundation. This work was supported in part by grant # UL1 TR000043 from the National Center for Advancing Translational Sciences (NCATS, National Institutes of Health (NIH) Clinical and Translational Science Award (CTSA) program and grant # R01 DC014247 from NIDCD (to L.B.V.). L.B.D. was supported by a Rockefeller University Women & Science Fellowship and by an APS Postdoctoral Fellowship in Biological Science from the American Philosophical Society. L.B.V. is an investigator of the Howard Hughes Medical Institute.

## AUTHOR CONTRIBUTIONS

L.B.D. carried out all experiments with the exception of the miniport small molecule screen in Figures 5 and 6, which were carried out by K.E.B. L.R.E. and J.F.G. helped to design and carry out the small molecule screen in Figure 5. L.B.D. and L.B.V. together conceived the study, designed the figures, and wrote the paper with input from all authors.

## MATERIALS AND METHODS

### Mosquito rearing and maintenance

*Aedes aegypti* wild-type laboratory strains (Orlando) were maintained and reared at 25 - 28°C, 70-80% relative humidity with a photoperiod of 14 hours light: 10 hours dark (lights on at 7 a.m.) as previously described (DeGennaro et al., 2013). Adult mosquitoes were provided constant access to 10% sucrose. Adult females were blood-fed on mice for stock maintenance, on human subjects for *NPYLR7* mutant generation, and on human subjects or sheep blood delivered via Glytube membrane feeders (Costa-da-Silva et al., 2013) for egg-laying and host-seeking experiments. Female mosquitoes were fasted for 14 - 24 hours in the presence of a water source prior to behavioral experiments. Blood-feeding procedures with live hosts were approved and monitored by The Rockefeller University Institutional Animal Care and Use Committee and Institutional Review Board, protocols 15772 and LV-0652, respectively. Human subjects gave their written informed consent to participate.

### *NPYLR7* mutant strain generation

The *NPYLR7* gene was mutated using CRISPR-Cas9 methods as previously described (Kistler et al., 2015). In brief, a 23 nucleotide guide RNA was designed to target the *NPYLR7* gene (target sequence with PAM underlined: ACCTCGATCGTCGGCAACG<u>TGG</u>). Purified guide RNA (40 ng/μl), Cas9 protein (300 μg) and a DNA plasmid containing a homologous recombination sequence including a fluorescent markers (200 ng/μl) were injected into 1069 pre-blastoderm stage *Ae. aegypti* embryos (Orlando strain) at the University of Maryland Insect Transformation Facility. 460 G0 animals survived, for a final hatch rate of 43%. G0 pupae were sexed and separated into male and female groups prior to eclosion. Male and female G0 adults were outcrossed to wild-type Orlando animals in batches of 20 G0 and 20 wild-type mates. F1 animals were screened for fluorescence to detect insertion of the fluorescent marker, but none were recovered. We therefore screened F1 animals for gene-disrupting insertions/deletions at the *NPYLR7* locus. 78 F1 animals were intercrossed and pooled into groups of 3 females and 3 males and analyzed with Illumina MiSeq for insertions/deletions surrounding the cut site. Animals were pooled into groups of 3 for genomic DNA extractions and MiSeq amplicon generation. PCR primers used to generate MiSeq amplicons (LD98MSF [TCGCCTCGCTCCGCTTCCAGA] and LD98MSR [CACCGAGATGGCCTGGGAGTA]). Genotypes were confirmed using Sanger DNA sequencing (Genewiz). Mutants were blood-fed on human subjects until a stable line was generated, and subsequently maintained by blood-feeding on mice.

### Peptide receptor cloning

*Ae. aegypti* NPYLR-expressing plasmids were previously described (Liesch et al., 2013). Full-length cDNAs for all other predicted peptide receptors in *Ae. aegypti* L3 and L5 genome annotations were cloned from cDNA isolated from 10 whole mosquitoes (Liverpool strain) or synthesized by GenScript and subcloned with XhoI-NotI into the pME18s vector for expression in mammalian cells (Supplemental Data File 1). For cases in which multiple isoforms were predicted from genome annotation, we tested the isoform generated from cDNA.

### Cell-based assays

HEK293T cells (ThermoFisher Scientific) were maintained using standard protocols in a Thermo Scientific FORMA Series II – Water Jacketed carbon dioxide incubator. Cells were transiently transfected with 1 μg each of plasmid expressing GCaMP6s (Chen et al., 2013), mouse Gqα15 (Offermanns and Simon, 1995), and a test receptor using Lipofectamine 2000 (Invitrogen). Transfected cells were seeded into 384 well plates (Greiner Bio-one), and incubated overnight in DMEM media (ThermoFisher Scientific) supplemented with Fetal Bovine Serum (Invitrogen) at 37**°**C and 5% carbon dioxide. Cells were imaged in reading buffer [Hanks’s Balanced Salt Solution (GIBCO) + 20 mM HEPES (Sigma-Aldrich)] using GFP-channel fluorescence of a Hamamatsu FDSS-6000 kinetic plate reader (Hamamatsu Photonics) at The Rockefeller University High-Throughput Screening and Spectroscopy Resource Center. Compounds were prepared at 3x concentration in reading buffer in a 384-well ligand plate (Greiner Bio-one). Plates were imaged every 1 sec for 5 min. 10 μl of compound was added to each well containing cells in 20 μl of reading buffer after 30 sec of baseline fluorescence recording. Fluorescence was normalized to baseline, and responses were calculated as either ΔF/F_0_ or as “max ratio” (maximum fluorescence level/baseline fluorescence level). Heatmaps were generated using heatmapper.ca (Babicki et al., 2016).

### *Ae. aegypti* peptide/receptor deorphanization screen

HEK293T cells were maintained, transfected, plated, and imaged as described above. *Ae. aegypti* peptides identified in previous peptidomics studies (Predel et al., 2010) or in collaboration with the Rockefeller University Proteomics Resource Center (Duvall et al., 2017) (Supplemental Data File 1) were synthesized by Bachem. Peptides were maintained as lyophilized powders or in 100% dimethyl sulfoxide (DMSO) (Sigma-Aldrich) stock solutions at −20**°**C. All peptides were tested at a 10 μM final concentration.

### High-throughput small molecule screen

HEK293T cells were maintained, transfected, plated, and imaged as described above with the exception that cells were plated and directly imaged in Fluorobrite DMEM media (ThermoFisher Scientific). Pilot experiments were performed using the LOPAC Library of Pharmacologically Active Compounds to optimize screening protocols and controls. Compound libraries maintained at the Rockefeller High-Throughput Screening and Spectroscopy Resource Center include: AMRI (50,000 compounds), AnalytiCon (700 compounds), BioFocus DPI (10,150 compounds), Chem-X-Infinity (4,000 compounds), ChemBridge (65,638 compounds), ChemDiv (126,000 compounds), Enamine (79,921 compounds), Edelris (2000 compounds), Greenpharma (240 compounds), Life Chemicals (30,272 compounds), SPECS (4051 compounds), Chiral Centers Diversity (3289 compounds), LOPAC1280™ (1280 compounds), MicroSource (2,000 compounds), Pharmakon (900 compounds), The Prestwick Chemical Library® (1120 compounds), NIH Clinical Collection (727 compounds), Tocriscreen Compounds (480 compounds), HTSRC Clinical Collection (303 compounds), Selleck Bioactive Compounds (1513 compounds). All compounds were prepared at 3x concentration in DMSO in reading buffer in a 384-well ligand plate and tested at a final concentration of 10 μM (with <1% DMSO final concentration). Responses were calculated as “max ratio” and each 384 well plate included a positive control column of 20 μM FMRFa3, an *Ae. aegypti* peptide agonist of NPYLR7, and a negative control column containing only reading buffer. 265,211 unique small molecule compounds were screened. Some compounds were present in multiple libraries and were therefore screened more than once. All screening data are available at PubChem (AID: 1259423).

The cutoff for cherry picking of compounds was set to Normalized Percent Activation (NPA) Max Ratio ≥ 40 and NPA Max-Min ≥ 40. This gave a total of 389 hit compounds (hit rate: 0.140%). Of these,13 compounds were removed from the list for reasons including abnormal cell density pattern, autofluorescence, or fluorescence signal spillover affecting neighboring wells. This resulted in a total of 376 compounds identified in the primary screen. Secondary screening included a counter-screen against non-transfected cells to exclude non-specific activation, and a 10-point dose-response curve starting at 20 μM with 1:2 dilutions.

### Fura-2 assay

HEK293T cells were maintained, transfected (without GCaMP6s), plated, and imaged as described above with the exception that cells were pre-loaded with the Fura-2 AM dye-loading solution provided with the Fura-2 Calcium Flux Assay Kit - No Wash, Ratiometric (Abcam) for 80 min at room temperature and calcium flux was ready using 340/380 nm excitation ratio channel of the Hamamatsu FDSS plate reader (Supplemental Data File 1).

### Glytube feeding

Females were fed sheep blood (Hemostat Laboratories), artificial blood, or saline using Glytube membrane feeders exactly as described (Costa-da-Silva et al., 2013). Artificial blood (Kogan, 1990) contained 15 mg/ml human γ-globulins, 8 mg/ml human hemoglobin, 102 mg/ml human albumin, 120 mM NaHCO_3_, and 1 mM ATP. Total protein content in blood is 208 - 230 mg/ml compared to 125 mg/ml in artificial blood (Kogan, 1990). The protein-free saline meal contained 120 mM NaHCO_3_ and 1mM ATP. For all drug-feeding experiments, compounds were diluted from stock solutions in 100% DMSO (Sigma-Aldrich), ensuring that the final concentration of DMSO was <1%. Meals were preheated for at least 15 min in a 42°C water bath and compounds and ATP were added to meals immediately before feeding and mixed by vortexing. Glytubes with 1.5 ml of each meal were placed on top of mesh on the mosquito cage, and females were allowed to feed through the mesh for at least 15 min. Fed females were scored by eye for engorgement of the abdomen and weighed to confirm feeding status. In the rare cases that females were scored as partially fed they were counted as non-fed and discarded.

### Egg-laying assays

7 to 14 day-old female mosquitoes were fed sheep blood using a Glytube membrane feeder (Costa-da-Silva et al., 2013). Immediately after confirmation of blood-feeding, individual females were placed in plastic *Drosophila* vials (25 mm diameter, 95 mm long) containing 5 ml of water and a 55 mm diameter Whatman filter paper (GE Healthcare) folded into a cone to act as an oviposition substrate. 5 days after the blood-meal, filter papers were removed, and eggs were manually counted.

### Response to mechanical stimulus

Startle responses were measured by loading individual 7 to 14 day-old females into LAM16 glass locomotor activity monitoring assay tubes (Trikinetics). Females were allowed to acclimate in the tubes for 30 min and then exposed to a mechanical stimulus in which the monitor was manually rotated 45 degrees and then returned to its initial position in a 3 sec pulse. Monitors were connected throughout the mechanical stimulation so that activity could be measured immediately after stimulus ceased. Activity counts were collected every 5 sec for 3 min. Animals were scored as activated if activity in the 10 - 40 sec post-stimulus window was significantly increased compared to the 30 sec pre-stimulus level using a paired t-test within each treatment group.

### Uniport olfactometer

Host-seeking behavior was measured using a uniport olfactometer exactly as described in (Liesch et al., 2013). Briefly, groups of 15 - 25 females, aged 7 - 14 days were loaded into small plastic canisters with mesh covering both openings obtained from the World Health Organization Vector Control Research Unit (Penang, Malaysia). Canisters were attached to a 1 m long plastic tube (19 cm diameter) that led to an attraction trap (14 cm long, 5 cm diameter), followed by a sealed chamber in which a human volunteer inserted a forearm. Humidified room air was carbon-filtered (Donaldson Ultrac-A) supplemented to a final concentration of 5% carbon dioxide using flow-meters (Cole Parmer). Mosquito-loaded canisters were attached to the olfactometer and given 5 min to acclimate prior to being released for a 5 min host-seeking trial. Mosquitoes were scored as attracted if they flew through the 1 m tube into the attraction trap within the trial period.

### Miniport olfactometer

Miniport olfactometers were constructed at the Rockefeller University Precision Fabrication Facility. Animals are loaded into 6” x 3” x 3” starting canisters in groups of 15 - 20. Humidified room air was carbon-filtered (Donaldson Ultrac-A) supplemented to a final concentration of 5% carbon dioxide using flow-meters (Cole Parmer). After a 5 min acclimation period, the canister was opened and animals were given access to an attraction trap containing one quarter of a nylon stocking previously worn by a human subject for 10 - 16 hours to collect odor. After a 10 min host-seeking period, the door between the start canister and attraction trap was closed and animals were scored as attracted if they flew into the attraction trap within the trial period. Detailed instructions for miniport olfactometer design, construction, and use can be found at https://github.com/VosshallLab/Miniport-Construction

### Blood-meal quantification

Female mosquitoes were starved overnight and fed sheep blood with 0.002% fluorescein (Sigma-Aldrich) using Glytube membrane feeders as described above. Females were sorted and stored at −20**°**C until fluorescence reading. Frozen mosquitoes were loaded into 1.5 ml Eppendorf tubes (ThermoFisher Scientific) containing 100 ml phosphate-buffered saline (PBS) (Lonza, BioWhittaker) and tissue was disrupted using a Kontes Pestle Pellet Grinder. Control wells containing 0, 0.5, 1.0, 2.0, 3.0, 4.0, or 5.0 μl of 0.002% fluorescein were combined with unfed control female mosquitoes to create a reference dilution curve. Homogenized plates were taken to the Rockefeller University High-Throughput Screening and Spectroscopy Resource Center, where 50 μl of homogenized solution from each well was transferred to a 384 well plate (Greiner Bio One, #784201) alongside 50 μl of the reference dilution curve in PBS. Fluorescent intensity for each well was measured using GFP-channel fluorescence of a Hamamatsu FDSS-6000 plate reader. Using the reference dilution curve, fluorescent measurements were converted back to volume (μl) of solution ingested.

### Mouse-in-cage assays

Females were fed sheep blood, saline, or saline with drugs using Glytube membrane feeders as described above. Fully engorged females were selected, weighed, and allowed to recover for 2 days. To distinguish between treatment groups, females were marked with colored powder (Slice of the Moon; Chameleon Colors) using the “shake in a bag” technique (Verhulst et al., 2013) and allowed to recover for 2 – 4 hours. Briefly, females were placed in mesh covered soup cups (Solo) and colored powder was applied using a 20 ml syringe (Becton Dickinson) inside a large plastic bag. Animals were then cold anesthetized and moved to a cage for recovery. Assays were performed in 30 cm^3^ screened mosquito cages (DP1000, MegaView). For each trial, groups of 18 - 24 females per treatment were given access to a single female BALB/c mouse (Harlan Laboratories) anaesthetized with ketamine:xylazine that was introduced into the center of the cage. Mosquitoes were given the opportunity to blood-feed for 15 min. The number of mosquitoes of each treatment group that blood-fed on the mouse was scored post hoc using a dissection scope (Nikon) and a LabCam iPhone adaptor (iDu Optics). Mosquitoes were scored as “freshly blood-fed” if there was visible engorgement with fresh blood in the midgut of if fresh blood was present after animals were crushed in a Kimwipe (Kimberly-Clark). A separate mouse was used in each experimental replicate.

## QUANTIFICATION AND STATISTICAL ANALYSIS

All statistical analysis was performed using Graphpad Prism Version 6. Data collected as percentage of total are shown as median with range. Data collected as raw values are shown as mean ± SEM or mean ± SD. Details of statistical methods are reported in the figure legends. Host-seeking behavior plots were generated using ggplot2 in RStudio 1.1.447, R 3.4.4 (R Core Team, 2017). The snake plot in Figure 4A was generated using Protter (Omasits et al., 2014).

## DATA AND SOFTWARE AVAILABILITY

All raw data in the paper are provided in Supplemental Data File 1. All plasmids described in this paper are available at Addgene (catalogue numbers: pending). Coding sequences for all peptide receptors described in this paper are available at Genbank (accession numbers: pending). All small molecule screening data are available at PubChem (AID: 1259423).

